# Evolutionary insights into an ancient fungal transition from land to sea

**DOI:** 10.64898/2026.06.22.733814

**Authors:** Kaylee E. Christensen, Aniliese Deal, Rachel B. Brem

## Abstract

Though fungi have largely been studied in the context of terrestrial niches, aquatic species that originated from terrestrial ancestors can be found across the fungal kingdom. To date, the mechanisms of these transitions from land to sea have remained poorly understood, and it is unclear what traits are associated with the specialization of a fungal to marine environments. Here we develop *Kluyveromyces* budding yeasts, sampled from terrestrial, estuarine, and marine niches, as a model for the evolutionary transition of fungi into the ocean. Comparative analyses of genomes from the genus revealed a contraction in genome size and gene number in aquatic *Kluyveromyces* compared to their terrestrial relatives. In laboratory culture, we uncovered evidence for phenotypic losses in aquatic *Kluyveromyces* species, namely desiccation and cold resistance relative to the terrestrial clade. Aquatic *Kluyveromyces* also exhibited better salt tolerance than terrestrial species, reflecting an evolutionary gain consonant with their provenance from seawater. Furthermore, in molecular-evolution analyses, we found robust signal for positive selection in the aquatic *Kluyveromyces* lineage, most notably at genes annotated for roles in oxidative stress. We interpret these results under a model in which the common ancestor of *Kluyveromyces* was terrestrial and first made a transition to estuaries and mangroves, leading to *K. siamensis* and *K. aestuarii*, which then primed further radiation into the marine niche, leading to *K. nonfermentans*. We propose that *Kluyveromyces* may be an ideal model to study these land-to-sea transitions.

## Introduction

Understanding trait heterogeneity in the natural world is a fundamental goal of evolutionary genetics. Radiation of species into new niches can be associated with the remodeling of life histories, including trait losses and gains. Tracing mechanisms of these events, especially when they happened millions of years in the past, remains a key challenge for the field. Fungi, with their small genomes and broad diversity, can serve as a powerful eukaryotic model for the discovery of principles that govern evolutionary innovation (Gostinčar et al., 2009; Naranjo-Ortiz & Gabaldón, 2020; Shang et al., 2016).

Evolutionary transitions between land and sea represent a major driver of diversification across the tree of life (Wei et al., 2026). In fungi, terrestrial lineages have, in many independent cases, given rise to marine descendants (Amend et al., 2019; Gladfelter et al., 2019; Raghukumar, 2017; Spatafora et al., 1998); indeed, phylogenetic analyses suggest that such transitions may be particularly frequent in fungi relative to other eukaryotic groups (Jamy et al., 2022). In addition to their potential as drivers of aquatic ecology (Breyer & Baltar, 2023; Seena et al., 2023; Sen et al., 2022), marine fungi are compelling candidate model systems for the study of the basic evolutionary biology of land-to-sea radiations. A number of phenotypes have been proposed as possible mechanisms for specialization by fungi to marine environments, including substrate preferences, adaptation to low-nutrient conditions, and halotolerance (Kurita et al., 2026; Lane et al., 2023). The latter, perhaps the most intuitive *a priori*, has not proven to distinguish marine from terrestrial fungal isolates in empirical work to date (Gladfelter et al., 2019); this would be expected if many fungi collected from ocean waters for such studies are generalist species washed in from land niches (Amend et al., 2019; Gladfelter et al., 2019; Kaewkrajay et al., 2020; Kohlmeyer & Kohlmeyer, 1979), manifesting an ancestral phenotype of salt resistance that is shared across the kingdom (Lages et al., 1999). Resolving the puzzle with analyses of salt tolerance in ocean-endemic fungi remains an ongoing need for the field. Furthermore, whether and how relaxed selection has shaped marine fungi—that is, the loss of traits that were under constraint in their terrestrial ancestors—has also been largely unexplored. Such losses could well include mechanisms among fungi to withstand instability in terrestrial environments (Bagamery et al., 2020), if abiotic stresses, *e.g.* local temperature (Valdes et al., 2021; Ximenes et al., 2016) and water availability, fluctuate less in marine settings. Genomic evidence for this notion has emerged from genome reductions in some marine fungi (Lane et al., 2023; Naranjo-Ortiz & Gabaldón, 2020; Stajich, 2017), analogous to the genomic streamlining known from comparisons of aquatic bacteria and archaea to their terrestrial relatives (Giovannoni et al., 2005; Gralka et al., 2023; Rodríguez-Gijón et al., 2022). As is the case with phenotypes, exactly which genome features are preferentially lost in marine fungi, and the evolutionary driving forces at play, have been less clear (though see (Lane et al., 2023)).

The genus of *Kluyveromyces* budding yeasts, which occupies a position flanked by terrestrial taxa in the fungal subphylum Saccharomycotina (James et al., 2020), encompasses clades associated with terrestrial and aquatic provenance. Best studied in the genus are the dairy yeasts *K. lactis* and *K. marxianus,* whose cultured isolates derive almost exclusively from terrestrial niches and have been the focus of an expanding evolution (Christensen et al., 2026; Friedrich et al., 2023; Varela et al., 2019) and biotechnology literature (Karim et al., 2020; Qiu et al., 2023; Spohner et al., 2016). These species form a putatively terrestrial clade with *K. dobzhanskii* and *K. wickerhamii,* which have likewise been defined on the basis of isolates from terrestrial settings (Belloch et al., 1997; Cai et al., 1996). Another clade of *Kluyveromyces* comprises isolates from saltwater and brackish water, hereafter referred to as the aquatic group: *K. aestuarii* and *K. siamensis* were named on the basis of isolates cultured from estuaries (Araujo & Hagler, 2011) and mangrove swamps (Am-In et al., 2008) respectively. To date, the cultured isolates for the third species in this clade, *K. nonfermentans,* are derived from marine sediments and from the surfaces of benthic invertebrate animals off the coast of Japan (Goshima, 2022; Nagahama et al., 1999). Given the ecologies of cultured *K. aestuarii*, *K. siamensis*, and *K. nonfermentans*, and their inferred evolutionary origin from a terrestrial ancestor, these species represent a compelling candidate case of an evolutionary transition from land to sea. The underlying mechanisms are unknown.

## Methods

### 1. Genomes and strains

#### 1.1 Gathering ecological data from GlobalFungi

We gathered ecological data for the tree figure from GlobalFungi (metadata downloaded via globalfungi.com on 2025-09-24) (Větrovský et al., 2020) using ITS sequences annotated as *Kluyveromyces* species (Table S1). *K. siamensis* and *K. wickerhamii* were not annotated in the GlobalFungi database and so data for these species are not shown. We gathered culture isolation data from CBS strains by the Westerdijk Fungal Biodiversity Institute (accessed via GBIF.org on 2025-09-24) (Verkleij, 2020) (Table S1). We further assigned Environment Ontology (ENVO) (http://www.ontobee.org/ontology/ENVO) biomes to each isolate based on given sampling information to match the terminology used by GlobalFungi.

#### 1.2 Genomes and strains used for analyses

Strains used for phenotyping are described in Table S2. For genomic analyses, the genomes used were as follows. For *K. marxianus* and *K. lactis* we used the reference genomes and annotations of Km31 and yLB72, respectively, from (Christensen et al., 2026). For *K. wickerhamii* we used the reference genome and annotation of UCD54-210 from (Sorrells et al., 2018). For *K. nonfermentans* strain NRRL Y-27343 (GenBank acc. no. GCA_030569915.1), the *K. aestuarii* strain NRRL YB-4510 (GenBank acc. no. GCA_003707555.2), and the *K. dobzhanskii* strain NRRL Y-1974 (GenBank acc. no. GCA_003705805.3) we used annotations from (Christensen et al., 2026). We generated a genome for *K. siamensis* strain CBS 10860 with the following steps: we grew an overnight culture in liquid yeast peptone dextrose (YPD), isolated DNA using the Quick-DNATM Fungal/Bacterial Miniprep Kit (Zymo Research), and sequenced ∼30M 150bp paired-end reads on an Illumina NovaSeq 6000 at the UC Berkeley Vincent J. Coates Genomic Sequencing Laboratory. We then assembled the genome with SPAdes v3.15.5 (Bankevich et al., 2012) using the default assembling mode, and generated an annotation with YGAP (accessed via http://wolfe.ucd.ie/annotation/ on 2025-09-13) which generates gene predictions based on annotations from similar yeast species (Proux-Wéra et al., 2012). As outgroups in phylogenetic analyses, we used genomes and annotations for the *Saccharomyces cerevisiae* strain S288C (GenBank acc. no. GCA_000146045.2), the *Eremothecium cymbalariae* strain DBVPG#7215 (ENA acc. no. GCA_000235365), the *E. gossypii* strain ATCC 10895 (GenBank acc. no. GCA_000091025.4), the *Lachancea nothofagi* strain CBS 11611 (ENA acc. no. GCA_900074755), and the *L. thermotolerans* strain CBS 6340 (GenBank acc. no. GCA_000142805.1).

#### 1.3 Genome quality assessment with QUAST

To ensure we were using high-quality genomes, we ran all *Kluyveromyces* genomes used through QUAST v5.2.0 without a reference genome (Gurevich et al., 2013). A summary table of the results is reported in Table S3.

#### 1.4 Proteome quality assessment with BUSCO

To ensure we were using high-quality proteomes and to account for annotation gaps, we ran all proteomes used through BUSCO v5.7.1 (Manni et al., 2021), which was installed via pip from the source due to conda dependency resolution issues. We used the ‘proteins’ mode and *S. cerevisiae* as the lineage dataset used (saccharomycetes_odb10 2024-01-08). A summary is shown in Table S3, as well as all of the genes that are missing or fragmented for *Kluyveromyces* species. A graphical summary of the results is reported in Figure S1.

### 2. Bioinformatics

#### 2.1 Orthologs

We input proteomes from the genomes and annotations from Section 1 for each species into OrthoFinder v2.5.5 (Emms & Kelly, 2019) to infer groups of genes of common evolutionary origin across species, resulting in 5,477 predicted orthogroups. From this set we earmarked the 2,917 one-to-one orthogroups with one copy in each species, including all *Kluyveromyces* and outgroup species, for molecular-evolution analyses (PAML and HyPhy; see sections 2.4 and 2.5); separately, we earmarked one-to-one orthogroups harnessing just *Kluyveromyces* species and *S. cerevisiae* for time tree analyses (see section 2.2). For gene family analyses (see section 2.6), we input all of the genes for each species from each gene family into Interpro (accessed 2026-07-31 InterPro 109.0) (Blum et al., 2021) to infer additional function. *S. cerevisiae* gene names were accessed from the Saccharyomyces Genome Database on 2026-09-26.

#### 2.2 Time tree construction

To construct a timed phylogeny of the *Kluyveromyces* genus for Figure 1A, we input aligned proteomes from each species, and *S. cerevisiae* as the outgroup, into the RelTime-ML function of MEGA-CC 11 (S. Kumar et al., 2012), using the tree topology from (Varela et al., 2019) with the placement of *K. siamensis* from (Am-In et al., 2008). We first generated proteomes using the genomes and annotations as in Section 1, then made protein-by-protein alignments with MUSCLE (Edgar, 2004) using the 3,330 one-to-one orthologs between *Kluyveromyces* species and *S. cerevisiae* called by Orthofinder as in Section 2.1. We concatenated alignments to generate one aligned proteomes file for input into RelTime-ML to generate the time tree. For exact time generations, we used the calibrations of: 27.4-28 mya between *K. marxianus* and *K. lactis*, 16.2-16.7 mya between *K. lactis* and *K. dobzhanskii*, and 51.1-51.2 mya between *K. marxianus* and *K. aestuarii* (Shen et al., 2018). The final RelTime-ML run used 1,685,729 sites with the Jones-Taylor-Thorton substitution model, clock type specific as local clocks, max relative rate ratio of 20, uniform rates among sites, and the “Use all sites” treatment of gaps/missing data. We formatted the final tree figure in iTOL (Letunic & Bork, 2024).

**Figure 1.**
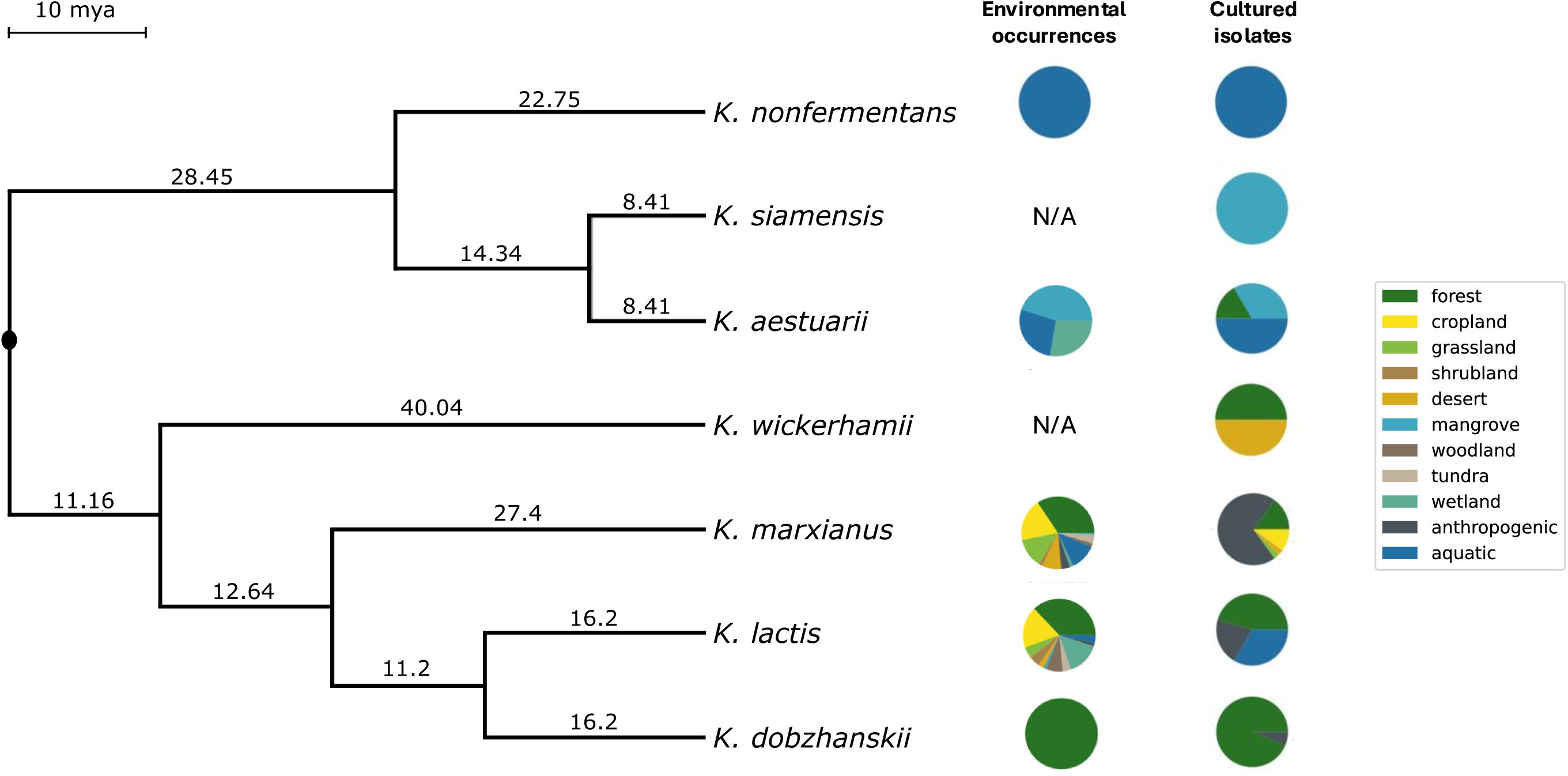
Ecology and phylogeny of the *Kluyveromyces* genus. Phylogenetic tree of *Kluyveromyces* species, scaled in millions of years (mya). Pie charts show relative occurrences of each species in different environments as detected from metabarcoding analyses in the GlobalFungi database (Environmental occurrences) and in the Centraalbureau voor Schimmelcultures collection (Cultured isolates).

#### 2.4 Finding evidence for positive selection (PAML)

As input to phylogenetic analyses of *Kluyveromyces* species and outgroups, we used genomes and annotations as in Section 1. For a given one-to-one orthogroup from Section 2 we generated protein alignments using MUSCLE v5.1 (Edgar, 2004). We then used this alignment and its nucleotide sequences as input into PAL2NAL v14.1 (Suyama et al., 2006) using the - nogap flag to remove gaps to generate a codon alignment. For each orthogroup in turn, we ran the PAML (Yang, 2007) codeml package run using the branch-site model (J. Zhang et al., 2005) to evaluate the fit to the sequence data of a model in which regions of the coding sequence are under positive selection for amino acid variation along a focal (foreground) branch and not on any other branch from the tree (the background branches). We used as input to PAML the codon alignment for the orthogroup and the topology of the phylogenetic tree from (Shen et al., 2018). The calculation proceeded in two steps. First, we fit a null model in which all branches of the tree had the same protein evolutionary rate (PAML codeml run with options verbose = 1, seqtype = 1, clock = 0, model = 2, NSsites = 2, fix_omega = 1, omega = 1), yielding a protein evolutionary rate and log-likelihood of the data under the model. Next, we re-fit using an alternative model in which the branch leading to the aquatic clade as the foreground (PAML codeml run with options verbose = 1, seqtype = 1, clock = 0, model = 2, NSsites = 2, fix_omega = 0). Outputs were the codon sites assigned to the class with an excess of amino acid changes in the aquatic clade relative to silent changes, and their posterior probabilities under Bayes Empirical Bayes inference (Yang et al., 2005); the codon sites assigned to classes with relaxed and purifying selection, respectively, along the background lineages, and their posterior probabilities; and the log-likelihood of the data under the model. The *p*-value assessing the improvement in fit to the data of the alternative model relative to the null was calculated from a likelihood ratio test (a χ^2^ test on twice the difference of likelihoods with one degree of freedom). *p*-values were corrected for multiple testing by the Benjamini/Hochberg method using the multipletests function from statsmodels (Seabold & Perktold, 2010). Separately, we repeated the branch-site test using the terrestrial *Kluyveromyces* lineage as the foreground. Results are listed in Table S5.

#### 2.5 Finding evidence for positive selection (HyPhy)

To complement our tests for positive selection using PAML in Section 2.3, we separately used the PAL2NAL alignments from section 2.3 as input for the aBSREL method in HyPhy (Pond et al., 2005). aBSREL uses a branch-site model similar to PAML to detect positive selection on individual branches. We ran aBSREL indicating either the aquatic ancestral or terrestrial ancestral branch of *Kluyveromyces* as the foreground. We collated the *p*-value as calculated by aBSREL for each orthogroup and corrected for multiple testing by the Benjamini/Hochberg method using the multipletests function from statsmodels (Seabold & Perktold, 2010). Results are listed in Table S5.

#### 2.6 Characterizing gene family evolution

We used the genomes and annotations as in Section 1 to generate a list of orthologs between the *Kluyveromyces* species and outgroups. Gene counts for orthogroups and a rooted phylogenetic tree, both outputs from OrthoFinder v2.5.5 (Emms & Kelly, 2019), were used as inputs for CAFE5 v5.1.0 (Mendes et al., 2021) to estimate gene family evolution across *Kluyveromyces* species and closely related outgroups. The base model was used and run twice to ensure convergence on the same lambda and final likelihood. Models with a different lambda for the aquatic clade were tested but were not predicted to have a different lambda from the rest of the tree, so the original model with one lambda was kept for further analysis. Gene families with zero gene copies at the root were excluded. A tree summarizing the increases and decreases for each taxon was made using CafePlotter v0.2.0 (https://github.com/moshi4/CafePlotter). Rapidly evolving gene families were identified based on having a significantly higher rate of evolution than the global mean rate estimated by CAFE5, with p-values corrected for multiple testing by the Benjamini/Hochberg method using the multipletests function from statsmodels (Seabold & Perktold, 2010). Gene families assessed to have significantly accelerated evolution (adjusted *p* < 0.05) at the node of the last common ancestor of the aquatic clade were reported in the heatmap in Figure 3C. Since the birth-death model used by CAFE5 is not reliable for small gene families (Hahn et al., 2005), we only reported results for the aquatic node where more than half of the species had a count of at least 1. To further account for this limitation, we developed a heuristic to find orthogroups in which the highest copy number for the aquatic *Kluyveromyces* species was lower than the lowest outgroup copy number, under the assumption that if all outgroups had a higher copy number then it was a loss in the aquatic clade. We excluded one result that we assumed to be essential, as it was RPS13, which BUSCO showed was missing in all three aquatic species annotations (Table S3); we concluded that this is likely an annotation issue rather than a true loss. The remaining eleven orthogroups found using the heuristic algorithm were plotted as a heatmap in Figure 3D.

### 3. Phenotyping

#### 3.1 Desiccation tolerance assay

For testing desiccation resistance differences between the aquatic and terrestrial species, we designed an assay for *Kluyveromyces* based on (Tapia et al., 2015). For each species, we streaked from a −80°C glycerol stock onto a yeast peptone dextrose (YPD) agar plate and incubated at 28°C for 2 days to achieve single colonies. We inoculated a single colony into 5mL of YPD media and then incubated the liquid culture overnight with shaking at 28°C. We then diluted an aliquot of this overnight culture into 10mL of YPD to achieve 0.05 OD_600_ and incubated the diluted culture with shaking at 28°C until it reached an OD_600_ of 0.2-0.4 (∼5-7 hours). This double-back dilution scheme ensures all cells are in exponential phase while undergoing the treatments. From the latter, for each replicate we sampled a volume of culture corresponding to 2 OD units, centrifuged, discarded the supernatant, resuspended in 1mL of phosphate-buffered saline (PBS), and again centrifuged and discarded the supernatant; we then incubated the pelleted cells in upside-down 2 mL Eppendorf tubes exposed to air in a 28°C incubator for two days or, for a control treatment, in closed tubes and incubated for two days at 28°C. After the two days, all liquid evaporated from the exposed tubes, providing the desiccation treatment, while the cell pellets in the closed tubes stayed hydrated with the leftover PBS. All species were assayed alongside each other, with each treatment replicated twice. We evaluated viability of the pelleted cells by estimating the total colony forming units (CFUs), where for each tube we resuspended the pellets in 1mL YPD, let sit for 10 minutes to rehydrate, formulated serial dilutions of this master culture in liquid YPD, spotted 3 µL of each dilution onto YPD agar plates, incubated the spotted plates for two days at 28°C, and counted the resulting colonies representing viable cells in the original pellets. Two-way ANOVA tests on the final CFU counts were performed using the anova_lm function from statsmodels (Seabold & Perktold, 2010) and reported in Table S2.

#### 3.2 Characterizing halotolerance and halophilicity

To assay growth in salt, we started with exponentially growing cells from a double back-dilution scheme as in Section 3.1. We made YPD media (Difco) supplemented with 3.5% or 7% NaCl (w/v) by adding 3.5g or 7g of NaCl (Fisher) per 100mLs of YPD, respectively. Since YPD has trace amounts of salts in unknown quantities, we reported the conditions based on the percentages of added salt. We took the second pre-culture at OD_600_ of 0.2-0.4 and inoculated it into 5 mL control YPD or added NaCl YPD at OD_600_ = 0.05, then incubated cultures with shaking at 28°C for ∼24 hours. We then recorded OD_600_ measurements and reported those values as total growth. Two-way ANOVA tests were performed using the anova_lm function from statsmodels (Seabold & Perktold, 2010) and reported in Table S2. Measurements were repeated in triplicate on different days.

#### 3.3 Characterizing growth in ecologically relevant temperatures

For growth in cold temperatures, we started with exponentially growing cells from a double back-dilution scheme as in Section 3.1 so that all cultures were in the same phase of growth when exposed to experimental conditions. We took the second pre-culture at OD_600_ of 0.2-0.4 and inoculated it into 5 mL YPD at OD_600_ = 0.05, then incubated cultures with shaking at 10°C for ∼72 hours to allow for detectable growth in the cold condition. We used 10°C as a representative midpoint of the range of temperatures of surface ocean water, which vary from ∼36°C to near-freezing depending largely on latitude. We then recorded OD_600_ measurements after growth in both temperatures and reported those values as total growth. Each treatment was repeated in triplicate. Two-way ANOVA tests were performed using the anova_lm function from statsmodels (Seabold & Perktold, 2010) and reported in Table S2.

#### 3.4 Measuring mitochondrial function with MTT

To measure mitochondrial activity, cells were cultured as previously described with the backdilution scheme to ensure that all were in exponential growth phase. We then carried out the MTT assay as previously described (Sánchez & Königsberg, 2006). Briefly, cells were harvested and the pellet resuspended in 100uL of MTT solution (100mM MES, 1M glucose, 5mg/mL 1-(4,5-Dimethylthiazol-2-yl)-3,5-diphenyltetrazolium Bromide (MTT)) and incubated for 1 hour at 28C. After incubation on ice for 5 minutes, cells were resuspended in DMSO, incubated at room temperature with consistent vortexing for 10 minutes, and then pelleted. The supernatant was then measured at 600nm in a spectrophotometer and diluted as needed if the readout exceeded the limits of the spectrophotometer. These final readouts were then reported as a bar chart (Figure S2A). The experiment was repeated twice.

#### 3.5 Measuring respiration and fermentation with Seahorse Xfe24 Analyzer

To measure metabolism more directly, a Seahorse Xfe24 Analyzer was used by adapting a previous protocol for yeast using a different Seahorse analyzer model (A. Kumar et al., 2024). Overnight cultures were grown in YPD to ensure robust growth. Since Seahorse medium is formulated for mammalian cells, we made synthetic defined medium designed for yeasts with similar specifications to Seahorse medium (YNB w/o amino acids (Difco), CSM (MP Biomedicals), 10mM glucose, 5mM HEPES, pH 7.0), and diluted the overnight cultures to OD600 = 0.05 in this medium. Cultures were grown until they reached OD600 of approximately 0.1-0.3 to ensure all cultures were in early exponential growth phase. 0.01OD cells in 500uL of synthetic defined medium were added to an Xfe24 culture plate that had been treated with 50ug/mL poly-D-lysine. The plate was spun at 500g for 5 minutes with no brakes to ensure an even coating of cells on the bottom of the plate. The plate was then incubated at 28C for approximately 45 minutes before being placed into the Seahorse analyzer with the temperature set at 28C. We reported the first three timepoints to ensure that oxygen levels were stable and the growth of the cells did not impact the measurements. There were 2-3 technical replicates for each species. The experiment was repeated twice. For extracellular acidification rate (ECAR), measurements were divided by the average of ECAR measurements for *K. marxianus* for the same day to account for variability; the same was done for oxygen consumption rate (OCR).

## Results

### Phylogeny and ecology of the *Kluyveromyces* genus

As a first window onto the evolutionary history of the *Kluyveromyces* genus, we elected to use a phylogenetic approach. We aligned the protein sequences of 3,330 gene orthologs between the *Kluyveromyces* species and *S. cerevisiae* and used them as input into a time-tree inference pipeline based on calibration with previously calculated divergence times (Shen et al., 2018).

The resulting model yielded an estimate for the timing of the split of the putative terrestrial and aquatic *Kluyveromyces* clades at ∼50 million years ago, and the divergence of the estuarine *K. aestuarii* and *K. siamensis* from the deep-sea *K. nonfermentans* at ∼23 million years ago (Figure 1).

We next sought an ecological evaluation of *Kluyveromyces* with respect to land and sea, in light of the convention in which a yeast is considered an endemic marine species when it is found exclusively from ocean sampling (Kohlmeyer & Kohlmeyer, 1979). Toward this end, to complement the literature focused on cultured isolates of *Kluyveromyces*, we surveyed metabarcoding data sources for each species in the genus in the GlobalFungi database (Větrovský et al., 2020). The results revealed recovery of *K. marxianus, K. lactis,* and *K. dobzhanskii* from primarily terrestrial sources, with a few reports from the ocean surface; *K. aestuarii* only from aquatic, wetland, and mangrove environments; and *K. nonfermentans* only from the ocean (Figure 1 and Table S1). *K. siamensis* and *K. wickerhamii* had no representation in the GlobalFungi resource. The latter notwithstanding, our results for all other *Kluyveromyces* species agreed with classic reports of the provenance of their cultured isolates (Figure 1).

Together, these data are consistent with the aquatic clade of *Kluyveromyces* as primarily occupying niches in estuaries or the ocean. Likewise, our data provide robust ecological support for the terrestrial *Kluyveromyces* clade as species found mostly on land; their occasional appearance on the ocean surface likely reflects cases of facultative migrants or their descendants that can occupy ocean environments but have not specialized to them ecologically, as is thought to be true for many other terrestrial yeast lineages (Kaewkrajay et al., 2020).

### Trait divergence between aquatic and terrestrial *Kluyveromyces*

Next, we sought phenotypes that distinguished the aquatic *Kluyyveromyces* from their terrestrial relatives, focusing on abiotic stress responses that could have potential relevance for aquatic life histories. We first hypothesized that desiccation tolerance would be under tighter constraint in terrestrial species, given their prevalence in dry environments such as deserts, tundras, and grasslands (Figure 1). We subjected a representative strain of each *Kluyveromyces* species to desiccation, then assayed viability by rehydrating the treated biomass and testing the capacity of cells in the sample to seed living colonies. In this scheme, aquatic species exhibited four orders of magnitude less viability after desiccation than species of the terrestrial clade (Figure 2A), as expected if desiccation tolerance were an ancestral phenotype lost as the clade evolved under the unique selective pressures from an ecology partly or fully underwater. In an ANOVA with treatment (control vs desiccated) and clade (aquatic versus terrestrial species) as factors, both were significant with *p* < 0.0005 as was the interaction between them.

**Figure 2.**
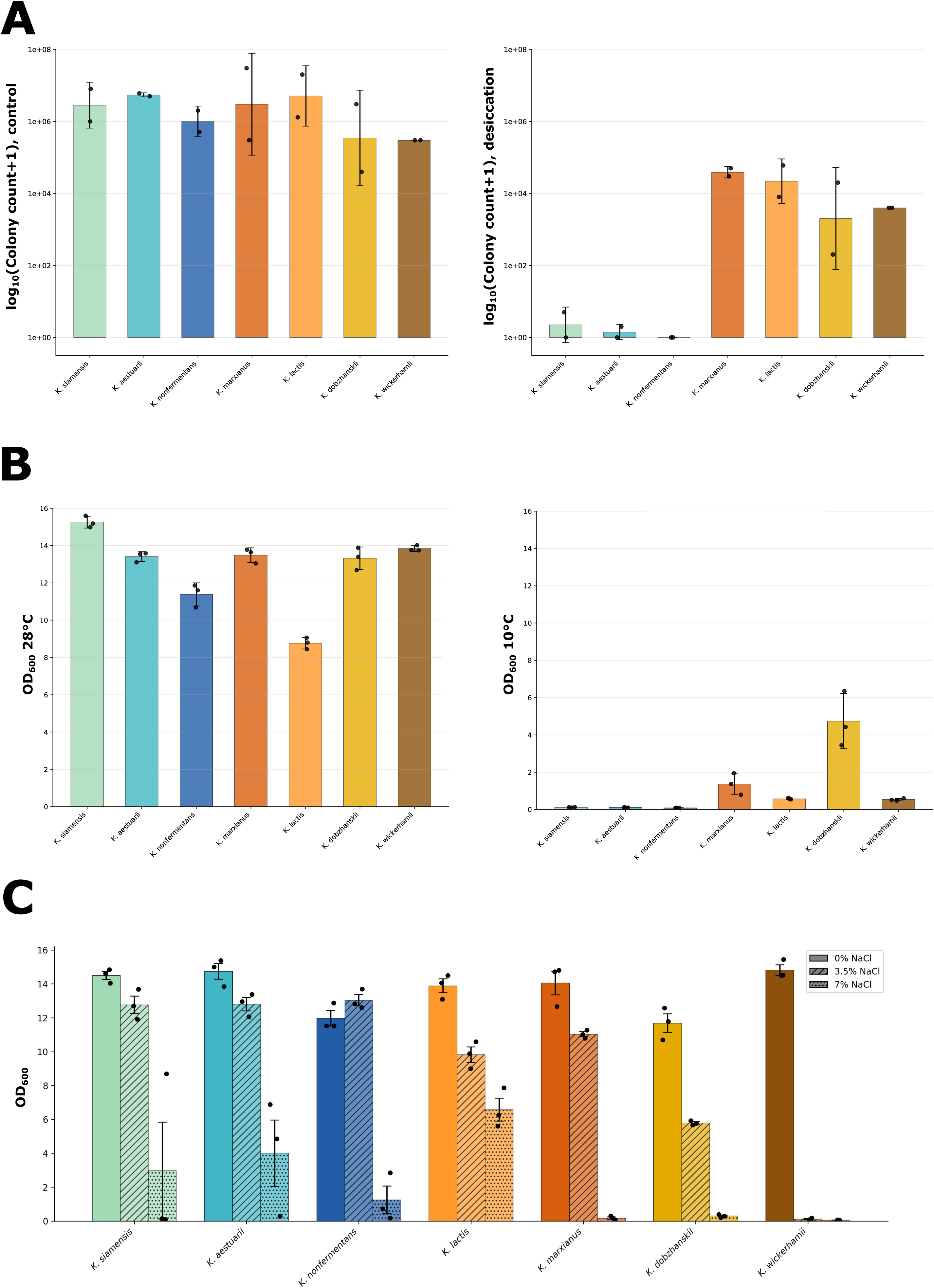
Trait divergence between aquatic and terrestrial *Kluyveromyces*. A) Desiccation survival. Each bar reports survival, in a culture of standardized input cell count, of the indicated *Kluyveromyces* species treated for 72 hours in desiccated conditions or a non-desiccated control. The *y*-axis reports the number of viable cells in the culture after treatment, as reported by colony growth from its inoculum when spotted onto agar plates; the bar height reports the mean and points report individual replicates. In an ANOVA with treatment (control vs desiccated) and clade (aquatic versus terrestrial species) as factors, both were significant with *p* < 0.0005 as was the interaction between them; see Table S2. B) Cold shock. Each bar reports biomass measured by OD_600_ of liquid culture in rich medium at 10°C or 28°C for 72 hours. In an ANOVA with treatment (10°C vs 28°C) and clade (aquatic versus terrestrial species) as factors, temperature was significant (*p* < 0.0005) as was the interaction term (*p* < 0.019); see Table S2. C) Salt tolerance. Each bar reports biomass measured by OD_600_ of liquid culture, after 24 hours of growth, of the indicated *Kluyveromyces* species in rich medium with 0%, 3.5% or 7% NaCl added. In an ANOVA with treatment (0% versus 3.5% NaCl) and clade (aquatic versus terrestrial species) as factors, both were significant with *p* < 0.0005 as was the interaction between them; see Table S2.

We further hypothesized that aquatic *Kluyveromyces* could experience more stable temperatures in their ecologies, and to explore this notion, we considered the response to temperature shock. As high-temperature growth has no clear ecological correlates across the genus (Christensen et al., 2026), we turned to a cold shock assay. In liquid growth assays upon a switch from 28°C to 10°C, all *Kluyveromyces* exhibited a defect relative to a constant-temperature control; this deleterious effect was more dramatic for aquatic species than for the terrestrial clade (interaction p < 0.019 in an ANOVA with treatment [10°C vs 28°C] and clade [aquatic versus terrestrial species] as factors; Figure 2B). Such a pattern is interpretable under a model of long-term evolution by aquatic *Kluyveromyces* under temperature conditions less subject to fluctuation than those on land, leading to a decline in the capacity to handle temperature change.

Next, we considered halotolerance, in light of the potential pervasive pressure on fungi to survive in the presence of salt in ocean water, estuaries, and mangroves. We used a standard liquid growth format to quantify biomass accumulation in salt medium of each species. In medium with 3.5% added NaCl, corresponding to the average salinity of seawater (Millero, 2010), all terrestrial species exhibited a drop-off in growth when compared to their performance in medium with no added salt (Figure 2C). By contrast, the deleterious effect of 3.5% added NaCl was more muted in the aquatic clade when the latter were analyzed as a group (Figure 2C; interaction *p* < 0.0005 from an ANOVA with treatment [0% versus 3.5% NaCl] and clade [aquatic versus terrestrial species] as factors). And the clade itself was not coherent with respect to salt response (interaction *p* = 0.005 from an ANOVA across aquatic species, with treatment [0% versus 3.5% NaCl] and species as factors), as the putatively marine species *K. nonfermentans* exhibited a growth advantage in 3.5% NaCl relative to the untreated control (Figure 2C). Thus *K. nonfermentans* thrives in, and estuarine species are uniquely tolerant of, salinity conditions associated with the ocean. All *Kluyveromyces* exhibited restricted growth at 7% NaCl (Figure 2C), arguing against extreme halophilicity in the genus.

### Genome features and gene loss in aquatic *Kluyveromyces*

A handful of studies have reported genomic streamlining and decreased GC content in marine fungal genomes (Lane et al., 2023; Naranjo-Ortiz & Gabaldón, 2020; Stajich, 2017), consistent with the extensive literature describing genomes of marine bacteria and archaea (Giovannoni et al., 2005; Gralka et al., 2023; Rodríguez-Gijón et al., 2022). We anticipated that similar trends would manifest in aquatic *Kluyveromyces* species. As a first test of this prediction, we analyzed *Kluyveromyces* genomes, finding with smaller genome sizes, lower GC content, and fewer genes in aquatic clade compared to terrestrial species (Figure 3A-B). Quality assessments ruled out artifacts from incomplete coverage as a driver of these effects (Table S3 and Figure S1).

**Figure 3.**
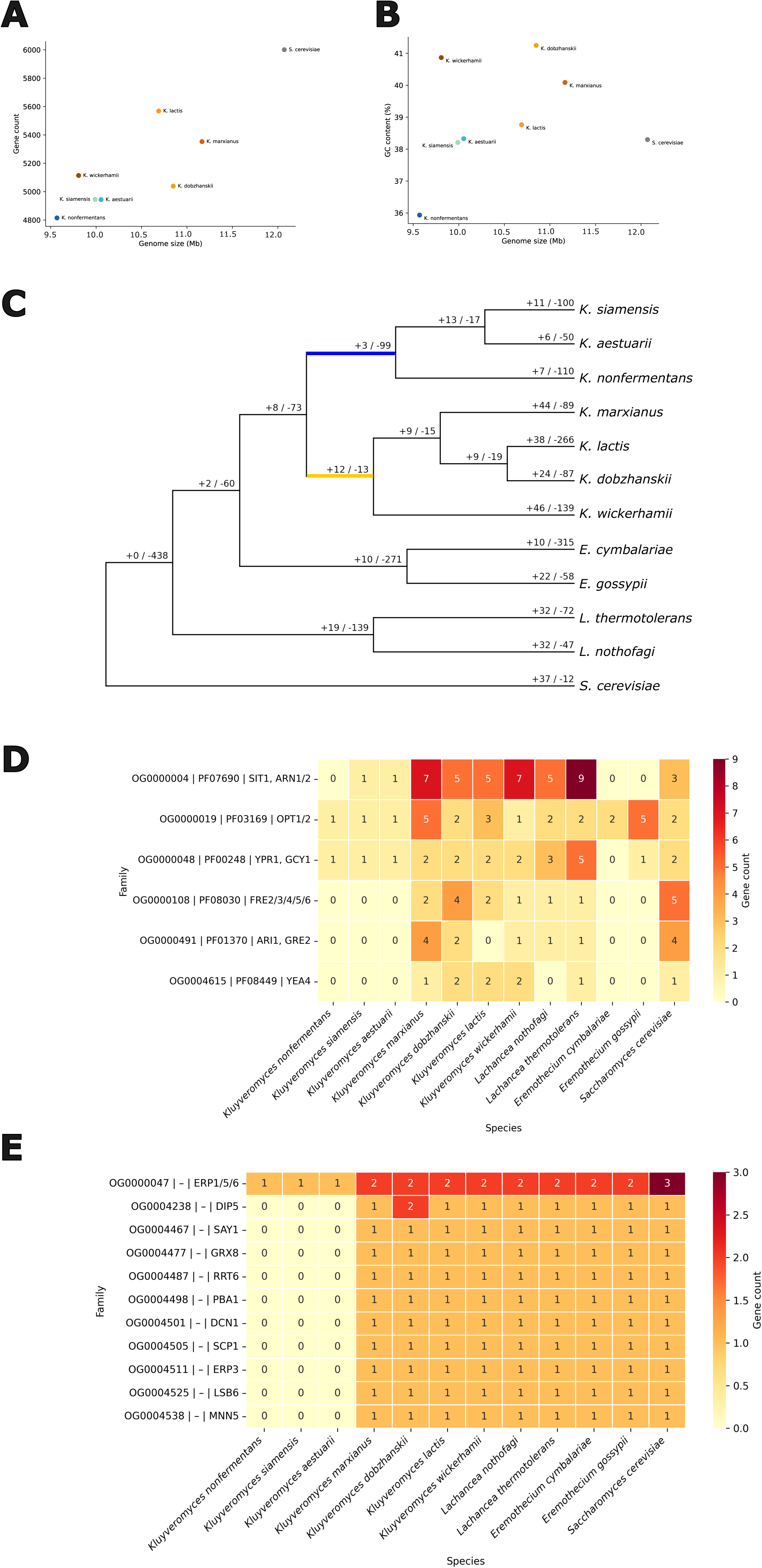
Genomic remodeling, gene loss, and relaxed selection in aquatic *Kluyveromyces*. A) Each point reports genome statistics for the indicated species. The *y*-axis reports genome size in megabases (Mb) and the *x*-axis reports the total number of annotated genes. B) Data are as in (A) except that the *y*-axis reports GC content (percentage of total base pairs). C) Summary of gains and losses across *Kluyveromyces* and outgroups. The blue branch represents the branch leading to the aquatic *Kluyveromyces* (i.e. the last common ancestor of the aquatic species), and the yellow branch represents the branch leading to the terrestrial *Kluyveromyces* (i.e. the last common ancestor of the terrestrial species). D) A heatmap of gains and losses for *Kluyveromyces* and outgroups for gene families that were rapidly contracting in the last common ancestor of the aquatic *Kluyveromyces*. To the left are labels reporting the gene family label from OrthoFinder, the most common Pfam ID associated with the genes in that gene family based on InterPro annotation, and the names of the orthologs from *S. cerevisiae.* E) A heatmap of gains and losses for *Kluyveromyces* and outgroups for gene families based on our heuristic algorithm. To the left are labels reporting the gene family label from OrthoFinder, the most common Pfam ID associated with the genes in that gene family based on InterPro annotation, and the names of the orthologs from *S. cerevisiae*.

Next, to shed light on gene losses harnessing a wider sampling of species, we constructed open reading frame alignments using sequences from *Kluyveromyces* and outgroup species, and we used them as input into orthogroup inference. We subjected the resulting orthogroups—each comprising genes with shared evolutionary ancestry, potentially including paralogs from a given species and orthologs across species—to tests for unique copy number change in aquatic *Kluyveromyces* as follows. We used CAFE5 (Mendes et al., 2021) to infer gene gains and losses across the phylogeny. Analyzing the complete set of such inferred changes, we detected an order of magnitude more inferred gene losses in the branch leading to the aquatic *Kluyveromyces* clade than along the terrestrial branch (99 versus 13 losses, respectively; Figure 3C). Next, focusing on large paralogous gene families, we identified those with evidence for accelerated rates of copy number evolution over and above the expectation under CAFE5’s null model. The results revealed accelerated loss rates in six orthogroups, including two annotated families of ion transporters and two families of NADPH-dependent reductases (Figure 3D and Table S4); interestingly, losses at all but two of these hits were also inferred in the non-fermenting terrestrial species *Eremothecium cymbalariae and Eremothecium gossypii* (Lachance et al., 2011) (Figure 3D and Table S4). As a complement, given the potential for errors in CAFE5 rate assessments for small gene families (Hahn et al., 2005), we developed a separate model-free heuristic method to test for orthogroups at which reduced copy number was shared across aquatic *Kluyveromyces*. This approach identified 11 orthogroups (Figure 3E and Table S4), including a transporter (Saccharomyces ortholog *DIP5*) and a glutathione-disulfide reductase (Saccharomyces *GRX8*). Together, these data underscore the prevalence of gene losses in aquatic *Kluyveromyces*, including in gene families relevant for redox chemistry and nutrient transport.

### Signatures of selection in aquatic *Kluyveromyces*

To deepen our understanding of the evolutionary forces acting on aquatic *Kluyveromyces*, we turned to a molecular-evolution approach. We used our whole-genus alignments of single-gene orthologs of *Kluyveromyces* and outgroup species as input into branch-site tests in the PAML (Yang, 2007) and HyPhy (Wertheim et al., 2015) packages to test each gene in turn for evidence of accelerated protein evolution on the branches leading to the aquatic and terrestrial clades, retaining genes that were top-scoring in analyses from both methods. The emerging set comprised 11 genes that had significant evidence for selection on the aquatic branch (Figure 4, Table 1, and Table S5). Six of these top hits were genes annotated with functions in mitochondria, including orthologs of the *Saccharomyces* genes *NDI1*, encoding the internal NADH-quinone oxidoreductase involved in mitochondrial electron transport; *KGD1*, encoding a component of the -ketoglutarate dehydrogenase complex; and *ATP1*, encoding a subunit of ATP synthase (Figure 4, Table 1, and Table S5). Assays of mitochondrial activity and oxygen usage yielded no evidence for divergent respiration phenotypes in aquatic *Kluyveromyces* (Figure S2A, S2B). Such a result argues against a role for adaptive amino acid variation in this facet of metabolism, though assays of extracellular acidification, a classic readout of yeast fermentation, were suggestive of a unique program in aquatic *Kluyveromyces* (Figure S2C). Together, these data attest to positive selection as a driving force for amino acid change in mitochondrial genes in aquatic *Kluyveromyces*, whose most likely phenotypic impact would be in aspects of function other than respiration *per se*.

**Figure 4.**
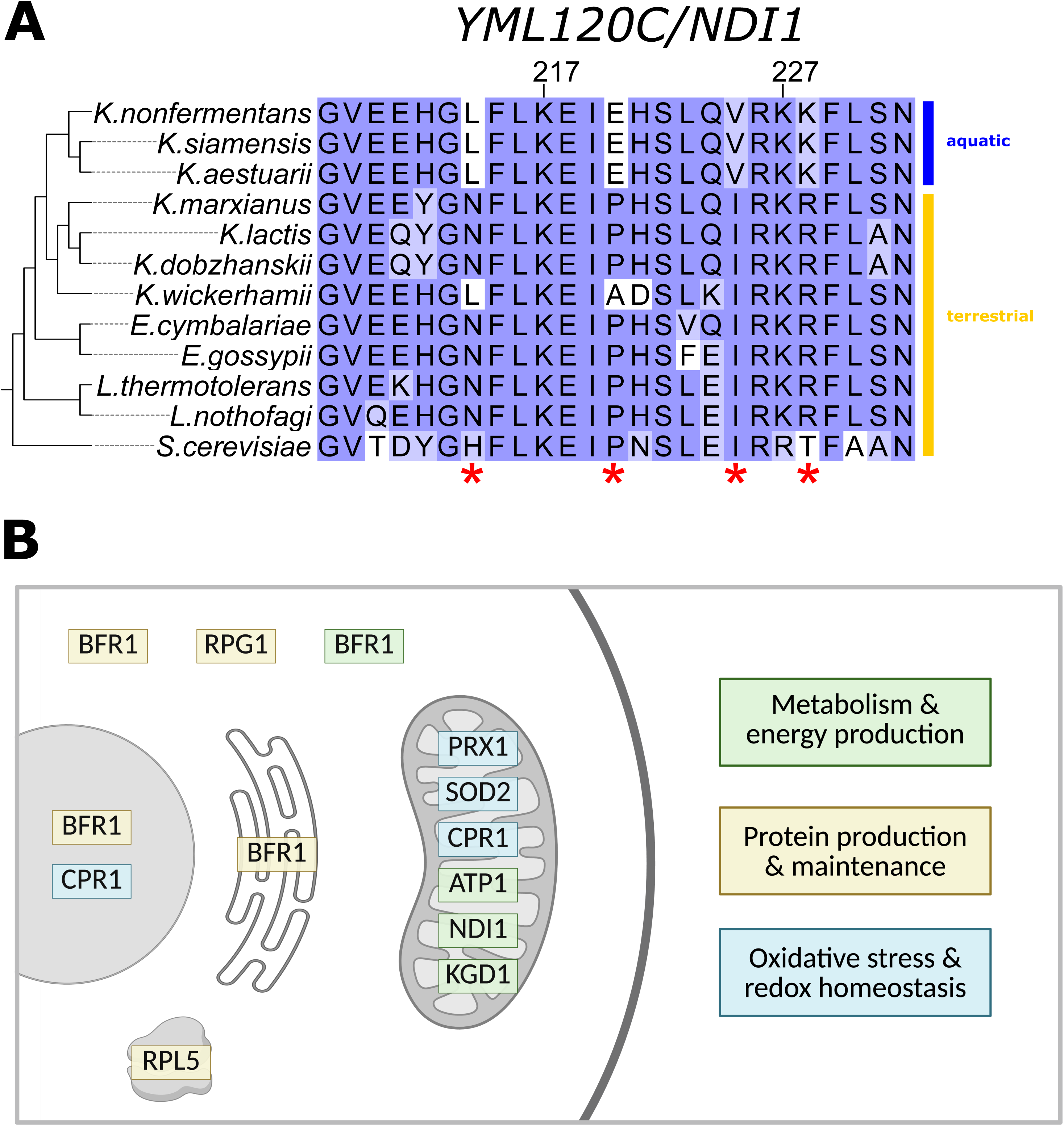
Positive selection in aquatic *Kluyveromyces*. A) An alignment of the orthogroup OG0000733 (containing orthologs of the *Saccharomyces* mitochondrial NADH dehydrogenase *YML120C/NDI1)*, an example case of positive selection in aquatic *Kluyveromyces*, with asterisks indicating sites with derived alleles in the aquatic clade (red dashed box). B) Results of a scan across orthogroups with HyPhy and PAML for evidence of positive selection in aquatic *Kluyveromyces* and where their encoded proteins are localized in the cell according to *Saccharomyces* annotations; the large grey circle denotes the nucleus, the open tubules denote the ER, the thick-walled grey organelle denotes the mitochondria, and the small irregular grey shape at bottom denotes the ribosome). Highlight color indicates general function: metabolism and energy production (green), protein production and maintenance (yellow), oxidative stress/redox homeostasis (cyan). Created with BioRender.com.

**Table 1.** Genes with evidence for unique positive selection acting on the branch leading to aquatic *Kluyveromyces*. Each row reports a gene with significant evidence for positive selection on the branch leading to the aquatic *Kluyveromyces* clade in both PAML branch-site and HyPhy aBSREL tests. For each gene the *K. marxianus* and *S. cerevisiae* orthologs are listed, along with the standard name and function according to the Saccharomyces Genome Database. Results from PAML and HyPhy on all tested orthologs are given in Table S5.

| *K. marxianus* gene | *S. cerevisiae* gene | Standard Name | Function |
| --- | --- | --- | --- |
| KM020F00410 | YOR198C | BFR1 | Component of mRNP complexes associated with polyribosomes; involved in localization of mRNAs to P bodies |
| KM020A05900 | YML120C | NDI1 | NADH:ubiquinone oxidoreductase |
| KM020H03660 | YBL064C | PRX1 | Mitochondrial peroxiredoxin with thioredoxin peroxidase activity; has a role in reduction of hydroperoxides |
| KM020C05480 | YBR079C | RPG1 | eIF3a subunit of the eukaryotic translation initiation factor 3 (eIF3) |
| KM020G01830 | YER065C | ICL1 | Isocitrate lyase; catalyzes the formation of succinate and glyoxylate from isocitrate, a key reaction of the glyoxylate cycle |
| KM020B05660 | YHR008C | SOD2 | Mitochondrial manganese superoxide dismutase; protects cells against oxygen toxicity and oxidative stress |
| KM020A01420 | Unknown | NA | Unknown |
| KM020F05080 | YDR155C | CPR1 | Cytoplasmic peptidyl-prolyl cis-trans isomerase (cyclophilin); catalyzes the cis-trans isomerization of peptide bonds N-terminal to proline residues |
| KM020E00290 | YIL125W | KGD1 | Subunit of the mitochondrial alpha-ketoglutarate dehydrogenase complex; catalyzes a key step in the tricarboxylic acid (TCA) cycle |
| KM020G01420 | YBL099W | ATP1 | Alpha subunit of the F1 sector of mitochondrial F1F0 ATP synthase |
| KM020B08140 | YPL131W | RPL5 | Ribosomal 60S subunit protein L5 |

## Discussion

Organismal lineages that have transitioned from land to sea represent a compelling study system for the dissection of how traits evolve. In fungi, understanding such transitions has remained a key challenge for the field, and insights into features that distinguish marine fungal species from their terrestrial counterparts have been at a premium for decades (Byrne & Gareth Jones, 1975; Damare et al., 2006; Gladfelter et al., 2019). Here, we have explored the *Kluyveromyces* genus as a model for the pursuit of mechanisms of fungi transitioning from terrestrial to aquatic environments. We have established *K. aestuarii*, *K. siamensis*, and *K. nonfermentans* as a clade that likely had terrestrial ancestors but whose extant members are consistently isolated from estuarine and marine niches. Over and above the previously reported carbon catabolism defects in *K. nonfermentans* (Nagahama et al., 1999), we have discovered additional phenotypes shared among aquatic *Kluyveromyces,* and we have also found evidence for pervasive gene loss as well as amino acid variants gained under positive selection in the genomes of the aquatic species.

In terms of specific phenotypes that we have found to distinguish aquatic *Kluyveromyces* from their terrestrial relatives, their advantage with respect to NaCl tolerance conforms to their inferred provenance in ocean and brackish water. This trait does not represent true halophilicity, *i.e.* peak performance under salt exposure, as in fungal halophiles recovered from salt pans (Gunde-Cimerman et al., 2009). Our result can be reconciled with the larger literature reporting little correlation between salt tolerance and marine ecology across fungi (Gladfelter et al., 2019; Kohlmeyer & Kohlmeyer, 1979) if the latter phenomenon were a consequence of experimental designs using wide fungal surveys, which include facultative marine species that have not made an evolutionary commitment to the seawater niche. That is, we propose that endemic marine fungi, including aquatic *Kluyveromyces*, could ultimately prove to be uniquely salt-tolerant. If so, we would expect such a phenotype to manifest with respect to other minerals in addition to the NaCl that is our focus here. Our work leaves open the genetic mechanisms underlying halotolerance in aquatic *Kluyveromyces*, as well as the unique sensitivity to desiccation and cold that we have observed in the clade. We expect that these traits might be highly genetically complex, as has been shown for species-specific thermotolerance in *Saccharomyces* (AlZaben et al., 2021; Walunjkar et al., 2025).

In light of our finding of amino acid variation with signatures of positive selection in mitochondrial genes of aquatic *Kluyveromyces*, but no divergence in respiration phenotypes, the adaptive benefit of such sequence changes remains unresolved. We note, however, that the affected loci are largely orthologs of oxidative stress protective factors (*PRX1, SOD2, CPR1*) or electron transport chain enzymes (*ATP1, NDI1, KGD1*) that represent a prime intracellular source of reactive oxygen species. As such, it is tempting to speculate that aquatic *Kluyveromyces* underwent evolutionary retooling of the proteins of these pathways in part to minimize redox stress. Such a logic could also explain the gene losses we noted in the clade in NADP(H)-dependent enzymes, if these contributed to a tuning of the NADP^+^/NADPH ratio to mitigate redox imbalance (Kwolek-Mirek et al., 2024). If aquatic *Kluyveromyces* had been under uniquely strong pressures to minimize reactive oxygen species production, and maximize oxidative protection, it could be a consequence of oxidative stress from exposure to salinity in the ocean niche, consonant with a previous study of the requirement for antioxidants in fungal salt adaptation (Ravindran et al., 2012). Additionally, ultraviolet radiation, temperature changes, or nutrient deficiency in marine environments could all also heighten the need for oxidative stress protection (Lesser, 2006; Xue et al., 2024).

In terms of metabolism proper, our finding of dampened extracellular acidification by aquatic *Kluyveromyces* relative to terrestrial species could be interpretable as a shared program of reduced fermentation in the former—with the most dramatic manifestation in *K. nonfermentans*, which lost the ability to ferment altogether (Lachance et al., 2011; Nagahama et al., 1999). If so, it would dovetail with the lost fermentation observed in other Saccharomycotina yeasts found in marine niches (Burgaud et al., 2011; Lachance et al., 2011; Mendonça-Hagler et al., 1985; Van Uden & Castelo-Branco, 1961). In aquatic *Kluyveromyces*, any such metabolic shift could be interrelated with the trend for paralog losses in gene families that we have found to be shared with non-fermenting *Eremothecium* species. Formally, however, reduced extracellular acidification by aquatic *Kluyveromyces* could also reflect changes in the production of organic acids, uptake of ammonium, or proton pumps (Liu et al., 2023; Peña et al., 1987; Schmidt et al., 2021); the latter could be an echo of the losses we have noted in transporter genes in aquatic *Kluyveromyces* genomes. Future work will be required to elucidate further whether the aquatic clade has undergone metabolic reprogramming and, if so, what evolutionary forces drove the changes.

Our discovery of small size and reduced GC content as features of aquatic *Kluyveromyces* genomes, especially in *K. nonfermentans*, echoes the results of previous genomic surveys of bacteria and archaea from aquatic environments (Giovannoni et al., 2005; Gralka et al., 2023; Rodríguez-Gijón et al., 2022). The small genomes of aquatic *Kluyveromyces* could have adaptive roles in promoting fast DNA replication, as has been hypothesized for other fungal clades (Lane et al., 2023; Naranjo-Ortiz & Gabaldón, 2020; Stajich, 2017). Additionally, both reduced genome size and low GC content could be an advantage to aquatic *Kluyveromyces* in the context of reduced access to nutrients needed for DNA synthesis, as suspected for marine bacteria and archaea (Gralka et al., 2023; Rodríguez-Gijón et al., 2022; H.-Y. Zhang et al., 2023). That said, many genome features in aquatic *Kluyveromyces* could also be attributable to a history of relaxed selection. Such effects could include a drop in GC content under relaxed selection for thermal stability (Yakovchuk et al., 2006) or DNA repair (Chen et al., 2004), and gene loss owing to reduced environmental variability in aquatic habitats or changes in carbon source specialization, if they eliminated the need for genes that were present in terrestrial ancestors, perhaps most saliently the transporters we have noted as lost in the aquatic clade.

In summary, our work has uncovered characters that are shared across aquatic *Kluyveromyces* (loss of desiccation and cold tolerance), as well as characters (genome size, GC content, salt tolerance) for which *K. nonfermentans* and terrestrial species represent the extremes of a continuum, with *K. siamensis* and *K. aestuarii* exhibiting intermediate values. Based on these findings, we propose a model in which the common ancestor of *Kluyveromyces* was terrestrial and that *K. aestuarii* and *K. siamensis* represent transitional states that ultimately led to the rise of *K. nonfermentans* as a marine specialist (Figure 5). In this scheme, in estuaries and mangroves, the common ancestor of the aquatic clade would have faced less-extreme versions of the challenges presented by life in the ocean; evolution in this environment would have primed descendants of this ancestor for further specialization to the ocean, leading to *K. nonfermentans.* As an alternative to the latter model, however, it remains possible that the features we have noted in aquatic *Kluyveromyces* arose in their ancestor independent of ecological pressures, given that phylogeny and environment are perfectly confounded in the genus. The future of the field will be devoted to pursuit of the mechanistic role of characters like those we study here, in the transitions that fungi have made into ocean niches.

**Figure 5.**
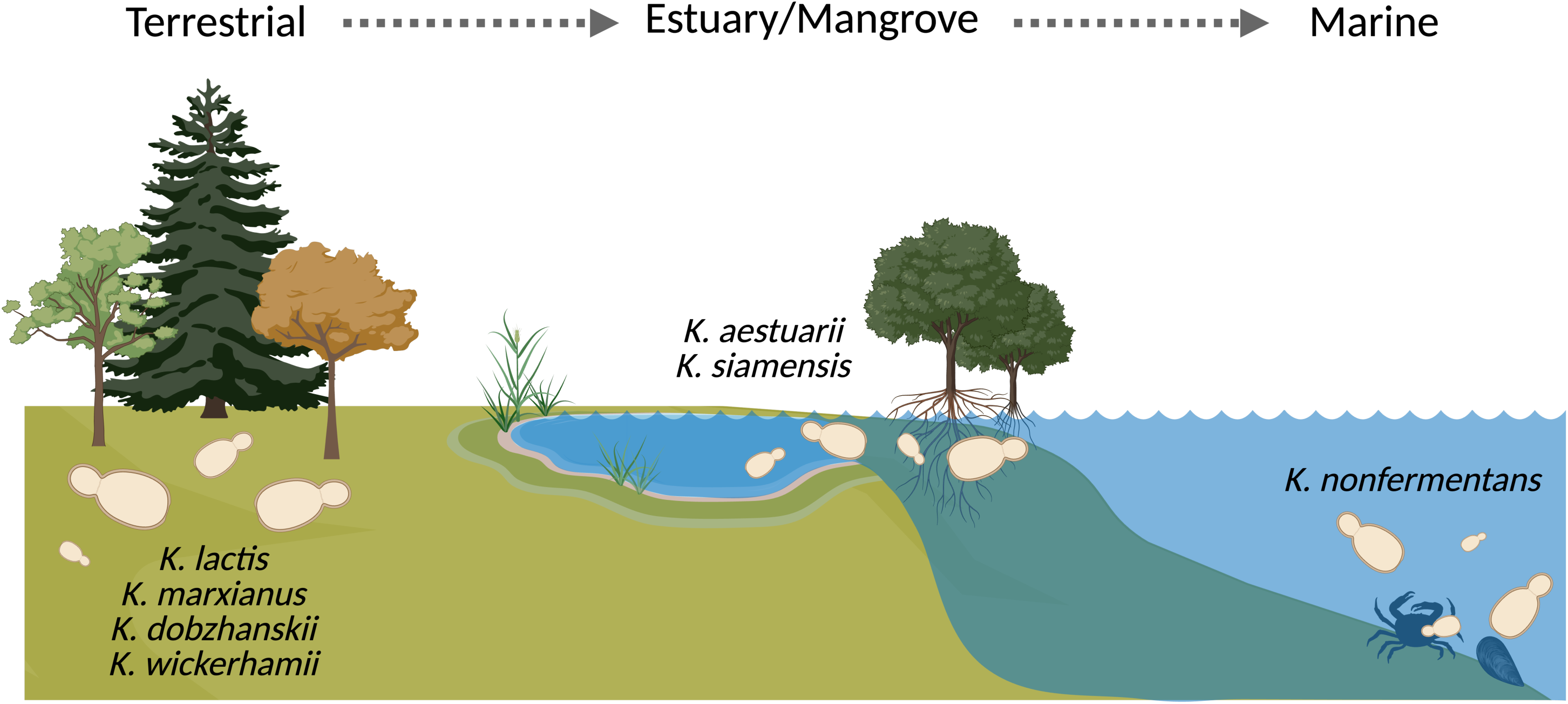
A model of the evolutionary trajectory of *Kluyveromyces* from land to sea. A proposed model of the evolutionary history of *Kluyveromyces* ecologies, in which migrants from an ancestral terrestrial population (extant species *K. dobzhanskii, K. lactis, K. marxianus, and K. wickerhamii*) gave rise to a mangrove/estuary population (extant species *K. aestuarii* and *K. siamensis*), from which migrants in turn gave rise to the extant marine species *K. nonfermentans*. Created with BioRender.com.

## Ethics statement

This study did not involve research on human participants, human data or tissue or vertebrate animals.

## Data availability statement

Genomic data and analysis scripts are available at https://github.com/kayleeec/aquatic_kluyv.git.

## Acknowledgements

We thank Anthony Amend and members of the Brem lab for helpful discussions. This work was supported by NIH 2R01GM120430 to R.B.B and stipend support for A.D. under NIH T32GM132022.

## Data Accessibility and Benefit-Sharing

Data accessibility: Raw sequence reads for *K. siamensis* are deposited in the SRA (BioProject PRJNA1472402). Genomes, proteomes, alignments, trees, and outputs from bioinformatics analyses are available at https://github.com/kayleeec/aquatic_kluyv Benefits generated: The data and results shared on public databases, as described above, ensure that the broader scientific community benefits from this research.

## Author Contributions

R.B.B. supervised the research and secured funding. K.E.C. carried out bioinformatics analyses and sequencing of *K. siamensis*. K.E.C. carried out the desiccation experiment. A.D. carried out genome and proteome quality analysis and growth in salt and cold temperature experiments, MTT and Seahorse experiments, CAFE5, and performed statistical tests on phenotyping experiments. A.D. drafted the manuscript with contributions from all authors. All authors reviewed and approved the final manuscript.

## Supplementary figure captions

**Figure S1.**
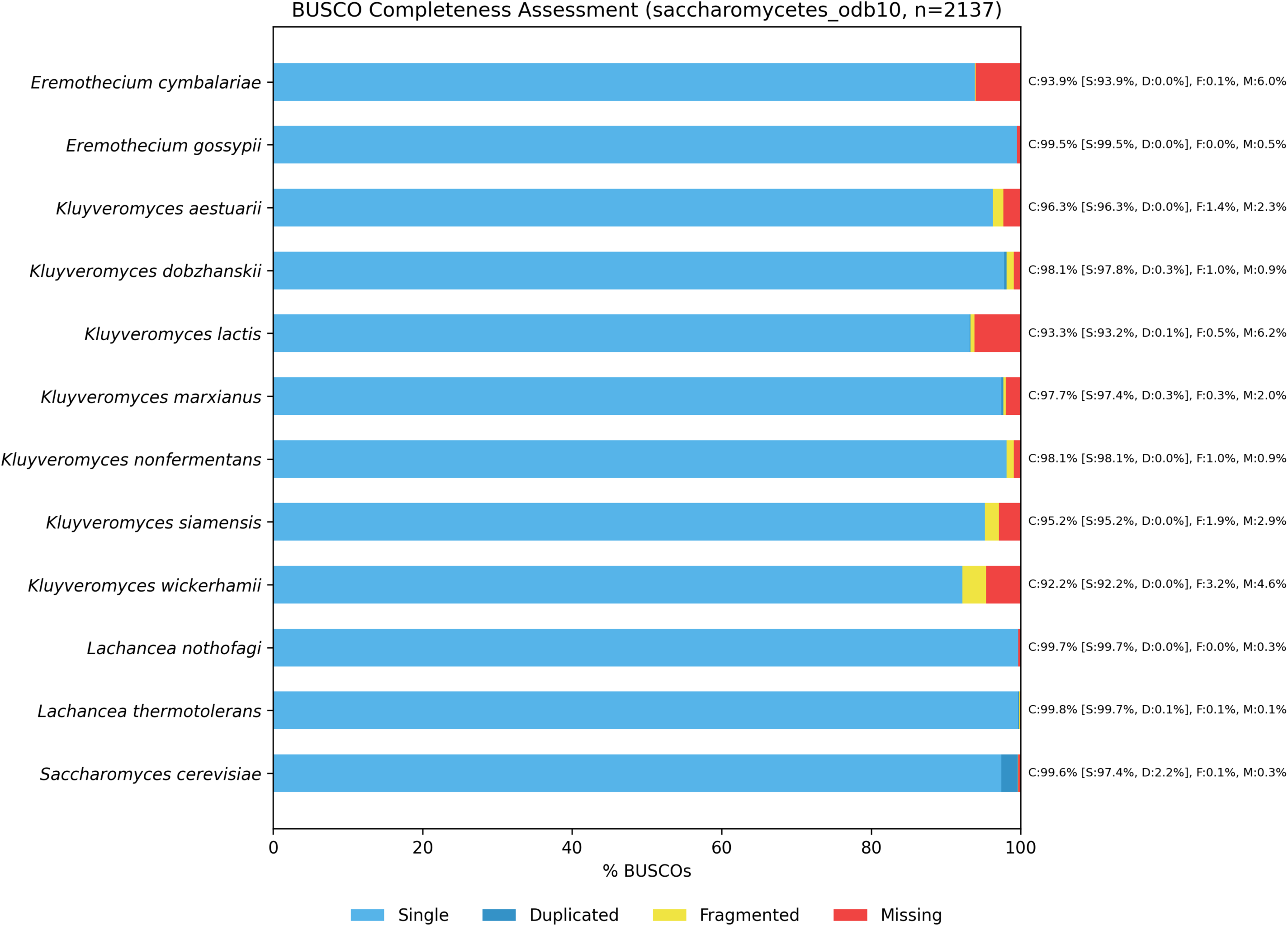
Proteome and annotation quality assessment with BUSCO. Species are listed in alphabetical order (left) with report results in BUSCO notation (right) in BUSCO notation (Complete [Single, Duplicated], Fragmented, Missing). Report results are visualized as stacked bar charts for each species.

**Figure S2.**
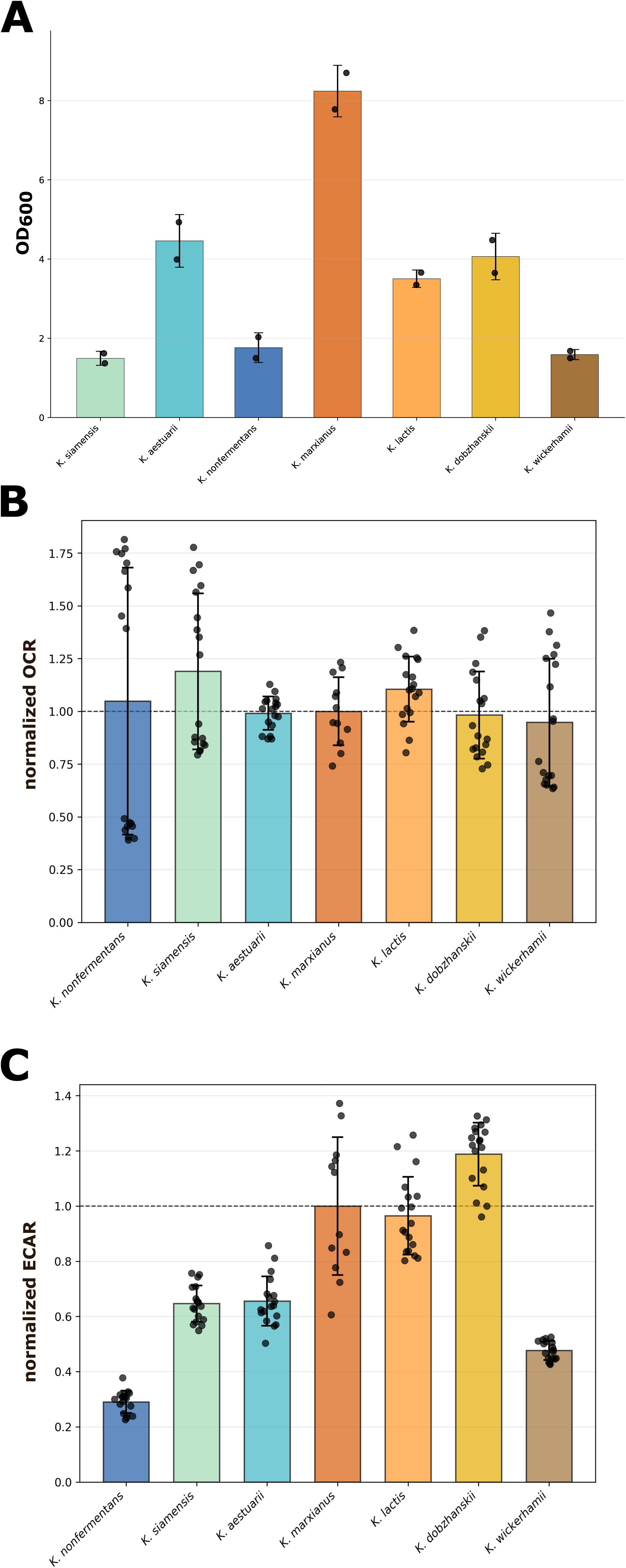
Profiling metabolic readouts across *Kluyveromyces*. A) Metabolic activity of mitochondrial enzymes as assayed by cleavage of MTT (3-[4, 5-dimethylthiazol-2-yl]-2,5-diphenyltetrazolium bromide). For each bar, the *y*-axis reports MTT cleavage by liquid cultures of the indicated *Kluyveromyces* species, as absorbance (optical density) at 600nm. The bar height reports the mean and points report individual replicates. B) Respiration as assayed by oxygen consumption rate (OCR), measured in pmol/min. For each bar, the *x*-axis reports OCR for the indicated species standardized to the analogous quantity from *K. marxianus*. C) Data are as in (B) except that the extracellular acidification rate (ECAR), measured in mpH/min, is reported.

